# Single Molecule Analysis of differential functional mechanisms of MtRecG while being exposed to variants of stalled replication-fork

**DOI:** 10.1101/2021.12.10.472094

**Authors:** Debolina Bandyopadhyay, Padmaja P Mishra

## Abstract

Helicases are motor proteins involved in multiple activities to carry out manipulation of the nucleic acids for efficient gene regulation. In case of roadblocks that can lead the replication machinery to get halted, a complex molecular surveillance system utilizing helicases as its key player ensures the halted fork to resume its duplication process. RecG, belonging to the category of Superfamily-2 plays a vital role in rescuing different kinds of stalled fork. Here, through adoption of single-molecule techniques we have attempted to probe the DNA unwinding features by RecG and tried to capture several stages of genetic rearrangement. An elevated processivity of RecG has been observed for the kinds of stalled fork where progression of lagging daughter strand is ahead than that of the leading strand. Through precise alteration of its function in terms of unwinding, depending upon the substrate DNA, RecG catalyzes the formation of Holliday junction from a stalled fork DNA. In summary, we have featured that RecG adopts asymmetric mode of locomotion to unwind the lagging daughter strand to facilitate Holliday junction creation which acts as a suitable intermediate for recombinational repair pathway.

## Introduction

In spite of being highly processive and inherently accurate, the genome duplication process in *Escherichia coli* or other prokaryotic cell is not intermittently completed without encountering impediments that can potentially pause the replisome unit.^1–3^ The progression of replication machinery is primarily hindered due to several considerable factors that includes single or double stranded breaks on the template, formation of unwanted secondary structures or binding of undesirable proteins to the DNA near the replication site. However, these impairments are resolved through well-coordinated damage specific repair mechanisms and that involves integrated action of various proteins especially helicases,^4^ playing critical roles in rescue of the challenged replication fork.^5–8^ The recombination specific RecBCD complex generates single stranded DNA through its nuclease/helicase activity and this is followed by strand invasion catalyzed by DNA strand exchange protein RecA.^9–11^ The nucleoprotein filament formed due to binding of RecA undergoes strand exchange with a homologous duplex giving rise to a Holliday junction, that in turn is resolved through RuvABC machinery.^12–15^ There are also certain kinds of lesions that bring replicating DNA unit carrying the infant strands in a mid-way halt, thus causing no net forward progression of the fork. Replication ‘fork reversal’ is a key protective mechanism that allows the forks to reverse their course when they encounter DNA lesions and resume DNA synthesis without chromosomal breakage.^16^ The process of fork reversal could either be spontaneous^16^ or catalyzed by protein that finally leads towards the advancement of the fork in a direction opposite to replisome movement.^17–19^

The RecG, in its monomeric form, is a key player in catalyzing the fork regression pathway and structurally modifies the stalled replication fork into Holliday junction through rearrangement of hydrogen bonds, thus making the halted fork a suitable target for recombination machinery.^20–22^ This works as a DNA translocase and unwinds a variety of branched DNA molecules in vitro, including Holliday junctions, D-loops, R-loops and various models of replication forks.^23,24^ It belongs to Superfamily 2 helicase (SFII helicase), the family of proteins possessing several other properties in addition to its classical helicase activity of unwinding the double-stranded DNA. RecG comprises of three structural domains, viz. domain I, II and III; the N-terminal domain I being the largest carrying more than half of the molecular weight of the protein.^25^ The ‘**wedge domain**’, a highly conserved ‘greek key motif’ is present in domain I, is linked to domain II and III via an alpha helical linker and the other two domains are linked through a cleft. The protein binds to the three-way junction through domain I, with the wedge domain clamping RecG tightly onto the DNA to facilitate splitting of the junction and stabilizing the DNA-protein complex.^26^ The domain II has ATP binding site and it possesses helicase activity in conjugation with domain III. Acting as an atypical helicase, the protein translocates in a direction opposite to the movement of replication machinery, that leads to splitting of the newly replicated strands from the template DNA.^25^ Being complementary to each other, the nascent leading and lagging strands anneal to form a four-way junction after getting separated from the respective parent strands. This four-way duplex intermediate structure is also famously known as ‘chicken foot intermediate’. Precise rearrangement of the chicken foot structure through branch migration leads to development of Holliday junction by making it a suitable resolvase substrate.^24,27^ It is not yet established and need an appropriate justification that how RecG is guided towards a stalled replication fork with an abundance as low as 7 molecules per cell. However, it has been hypothesized that within the milieu of genome, single stranded binding protein SSB acts as a guiding unit to locate RecG at the site of challenged replication fork for its rescue. ^28,29^ Since, other than binding to single stranded DNA and protecting them from degradation, SSB also interacts with a group of proteins for example PriA, RecQ, RecG and Topoisomerase III, called ‘SSB-interactome’ facilitating their actions in genome modifications. Once loaded, the fork reversal activity of RecG has been hypothesized over the years through several bulk assays by determining the affinity and unwinding activity of RecG for various substrates resembling replication halted structures.^21,30^ However, convincing experimental evidence in terms of translocation of the protein and mechanical quantitative insights into the detailed dynamics of the substrate DNA still remains elusive. And primarily, till date there is no direct observation for the motility of RecG in reverse direction promoting fork-reversal. It has also been found that RecG acts as a ‘stringent regulator’ for the rescue of stalled replication complexes and is capable of binding to variety of fork structures with a difference in affinity and functional aspects. Acting as a helicase with 3’-5’ polarity, other than branched DNA substrates RecG has also been found to process negatively supercoiled DNA ((-)scDNA), single-stranded DNA (ssDNA) or SSB coated M13 DNA. And, interestingly it has been verified that RecG has a stronger preference for (-)scDNA suggesting the phenomenon that substrate DNA should be converted from (+) to (-)scDNA^31^ for allowing RecG to function. Now, through deep functional analysis of every molecule via implementation of the high level of sensitivity and spatiotemporal resolution of the smFRET methods^32,33^, we have closely monitored the action and quantified the dynamical heterogeneity of every possible arrested replication forked DNA upon being allowed to bind with RecG. Taking the same substrate fork and by labelling, it with donor and acceptor at different positions we could monitor the helicase action of RecG in both forward and reverse direction separately. By amalgamation of all the results we come up with a final model of ‘asymmetric fork regression’ where selectively lagging strand unwinding is catalysed by RecG however processivity along the leading daughter strand is significantly less.

## Materials and Methods

### Purification of MtRecG (*Microbacterium tuberculosis* MtRecG)

MtRecG, cloned in pET28a (+) E. Coli expression vector (generous gift from Dr Ganesh Nagaraju) has been transformed in E. Coli BL21(DE3) cells.^30^ The transformed BL21(DE3) cells are grown in LB broth containing 50 µgml^-1^ Kanamycin. Once OD_600_ reaches 0.6, the cells are induced with 2 mM isopropyl-1-thio-β-D-galactopyranoside (IPTG) and allowed to grow overnight, specifically 16 hours at 18°C. The cells are then harvested by centrifugation at 5,000 rpm for 10 minutes at 4°C. Resuspension of the cell pellet is done in Lysis Buffer having the composition of 20 mM Tris-Cl (pH 8.0), 500 mM NaCl, 0.1 mM EDTA, 5 mM β-mercaptoethanol (β-ME), 10% glycerol. The resuspended cells are then treated with protease inhibitor cocktail (PIC purchased from Sigma) at a dilution ratio of 1:1000. Post resuspension the cells are subjected to sonication (12 cycles of 30 sec pulse and 30 sec gap) and then centrifuged at 10,000 rpm for 40 minutes keeping the temperature at 4°C. The supernatant collected after centrifugation are loaded in a column with Ni-NTA Agarose beads pre-equilibrated with lysis buffer. To get rid of non-specific proteins bound with the beads, the column is washed with 15 CV (Column Volume) Wash Buffer (20 mM Tris-Cl, pH8.0, 500 mM NaCl, 0.1 mM EDTA, 10% glycerol, 5 mM β-mercaptoethanol (β-ME) and 50 mM Imidazole), and the protein (MtRecG) is eluted in 1 ml fractions using Elution Buffer (20 mM Tris-Cl, pH 8.0, 500 mM NaCl, 0.1 mM EDTA, 10% glycerol, β-mercaptoethanol (β-ME) and 250 mM Imidazole). MtRecG, has the molecular weight of 80 kDa. The eluted samples are checked in polyacrylamide Gel Electrophoresis (SDS PAGE, 5% stacking and 10% resolving) and elute fractions with a single band corresponding to 80 kDa molecular weight are pooled down and subjected for buffer exchange against Dialysis Buffer (20 mM Tris-Cl, pH – 8.0, 500 mM NaCl, 5 mM β-mercaptoethanol (β-ME) and 10% glycerol) using Amicon Ultracentrifugal Filters having the cut-off of 50 kDa molecular weight. The centrifugation during buffer exchange has been done at 2500 rpm keeping the temperature is fixed at 4°C. Further, MtRecG is concentrated and the final concentration is determined using Absorbance at 280 nm, A_280_ value (Nanodrop Spectrophotometer) and dividing it with the ε-value of MtRecG (50,700 M^- 1^cm^-1^). The concentrated samples are stored in 4°C after a final check in SDS PAGE with 5% stacking and 10% resolving (Supplementary Figure S1), in 50 µl aliquots and used for smFRET experiment within 3-4 weeks of purification. The dilution of the protein is carried out in Reaction Buffer (50 mM Tris-Cl, pH 8.0, 50 mM NaCl and 5 mM MgCl_2_). All the buffers are filtered with 0.22 µm membrane filters before use.

### DNA Constructs

All the DNA constructs modified or unmodified used in this study are HPLC Purified and purchased from IDT (Integrated DNA Technologies) and used directly without any further purification. The DNA samples used for smFRET experiments are conjugated with a biotin labelled DNA strand to ensure immobilization via binding with streptavidin. The DNA sequences are modified with Cyanine 3 (Cy3) and Cyanine 5 (Cy5) at required positions (as explained in Supplementary Table 1), acting as donor and acceptor respectively in this study. All the DNA sequences used to make the desired substrate Fork structures are shown in Supplementary Figure S2. For annealing, the DNA strands are mixed together to make the final concentration to 20 µM. the fluorophores and biotin carrying sequences, (Cy3:Cy5:Biotin Strand) are mixed in the ratio of 1:1:1. The samples are first heated to 95^°^C and then slowly cooled down to 8^°^C for 2.5 hours and stored at -20^°^C till further experimental use. The 20 µM DNA stock samples are then serially diluted in Reaction Buffer to 100 pM typically termed as single molecule concentration, for exhibition of smFRET experiments.

### smFRET Data Acquisition and Imaging

All smFRET experiments are executed in a home-built prism-type total internal reflection microscopic^34^ (p-TIRFM) set-up explained in details elsewhere.^35,36^ The fluorophore labelled DNA used in this study are immobilized on pre-drilled quartz slides having microfluidic sample chambers made via attaching glass coverslips on the slides using double-sided tapes. The pre-processing of the quartz slides in order to make them ready for smFRET experiments has been explained elsewhere. The donor fluorophore, Cy3 is excited using 532 nm diode green laser (Laser Quantum, UK) and depending upon the position of the acceptor (Cy5) from the donor, the acceptor also gets excited through Forster’s Resonance Energy Transfer (FRET).^37^ The monochromatic laser beam is initially expanded to four times of it’s width followed by constricting it using a biconvex lens of focal length 100 mm. The ray is exposed on the sample in such a way, so that the incident angle is greater than the critical angle (61.5°) causing total internal reflection (TIR) and generation of evanescent waves, that excites the donor. The emitted signals from both the donor and acceptor are captured and separated using a 640 nm DCXR, dichroic mirror (Chroma) followed by projection on EMCCD (Electron Multiplying Charge-Coupled Device) camera. The movies are recorded using frame integration time of 35 ms. The data is processed using IDL script and single molecules with anticorrelated intensity of the donor and acceptor, single step photobleaching and having no blinking events are selected for further analysis.^38^ Every single molecule trace displays the intensities of Cy3 and Cy5 as a function of time and corresponding FRET Efficiency (E_FRET_) value is found through the ratio, I_A_/(I_A_ + I_D_). Data analysis has been carried out for ∼200 molecules through ebFRET and other self-written Matlab scripts are used for extracting the dwell time at a particular FRET state, FRET Efficiency histograms of individual molecules and transition density plots. Other than the compiled FRET Efficiency histogram, that carries ensemble information from all the molecules, we have generated superimposed FRET Efficiency histograms of all the individual molecules in order to get the account of every transition within all the molecules separately. All the individual raw FRET Efficiency versus time traces are idealized by fitting with Hidden Markov Model (HMM) Analysis^39,40^ (Viterbi Algorithm) through ebFRET^41^ interface and information generated regarding the variable states are used to extract the dwell time from all the molecules at a particular conformational state. The heat map of transition density plot (TDP) depicts the initial and final state during transition between two different fitted FRET states within the same molecule thus providing us quantitative information about transition dynamics.

The smFRET experiments of DNA-protein interaction have been carried out in Reaction Buffer. The DNA substrates are immobilized on streptavidin (0.2 mg/ml) treated fluid chambers and allowed to get incubated for 5 minutes. MtRecG diluted in reaction buffer is then flushed into the chamber. The reaction buffer, supplemented with oxygen scavenging system consisting glucose oxidase (2 mg/ml), Catalase (0.03 mg/ml) and glucose (0.4% by mass) as substrate, is flown into the chamber followed by acquisition of movies.^42^ Concentration of MtRecG has been kept constant to 500 nM throughout all the experiments. Also, 1mM Trolox is added to prevent photo blinking. All the experiments have been carried out at room temperature.

### smPIFE assays

In order to execute single molecule protein induced fluorescence enhancement (smPIFE)^43^, experiments, the desired fork DNA substrates (Supplementary Figure S3) are labelled with only donor at the junction. The dilution of the fork substrates was carried out in reaction buffer and they were immobilized on the slide along with oxygen scavenging system. The protein MtRecG diluted in Reaction Buffer (keeping the final concentration as 500 nM) and supplemented with oxygen scavenging reagents was stored in the reservoir made at one end of the chamber and during acquisition of the movie, after a certain fixed number of frames the protein was allowed to flow inside the chamber through suction using a silicon tube fixed at the other end acting as outlet.

## Results

### Repetitive unwinding by MtRecG at the junction of Open Forked DNA

Over the past several years MtRecG has been known as a unique protein that plays a vital role in reversing the direction of replication fork. In this study, we have tried to explore and quantify the differential helicase activity of MtRecG in forward and reverse direction on the types of forked DNA substrates those act as potential intermediates of a stalled replication process. To begin with, the helicase activity of MtRecG, typically the unwinding action have been examined right at the single strand-double strand (ss/ds) junction towards the duplex stem region of an open fork DNA, having 25 base pairs at the duplex region and both the arms flanking due to 10 nucleotide non-complementary single stranded overhangs (Supplementary Figure S2a). The binding of MtRecG at the ss/ds junction of an open fork is confirmed through smPIFE (Figure 1a). The junction was labeled with Cy3 moiety and once MtRecG is flown into the system, we found an approximate two-fold increase in Cy3 intensity within 2 seconds of introduction MtRecG into the chamber (Figure 1a). The enhancement in signal confirms binding of MtRecG in close proximity of Cy3 unit.

**Figure 1.**
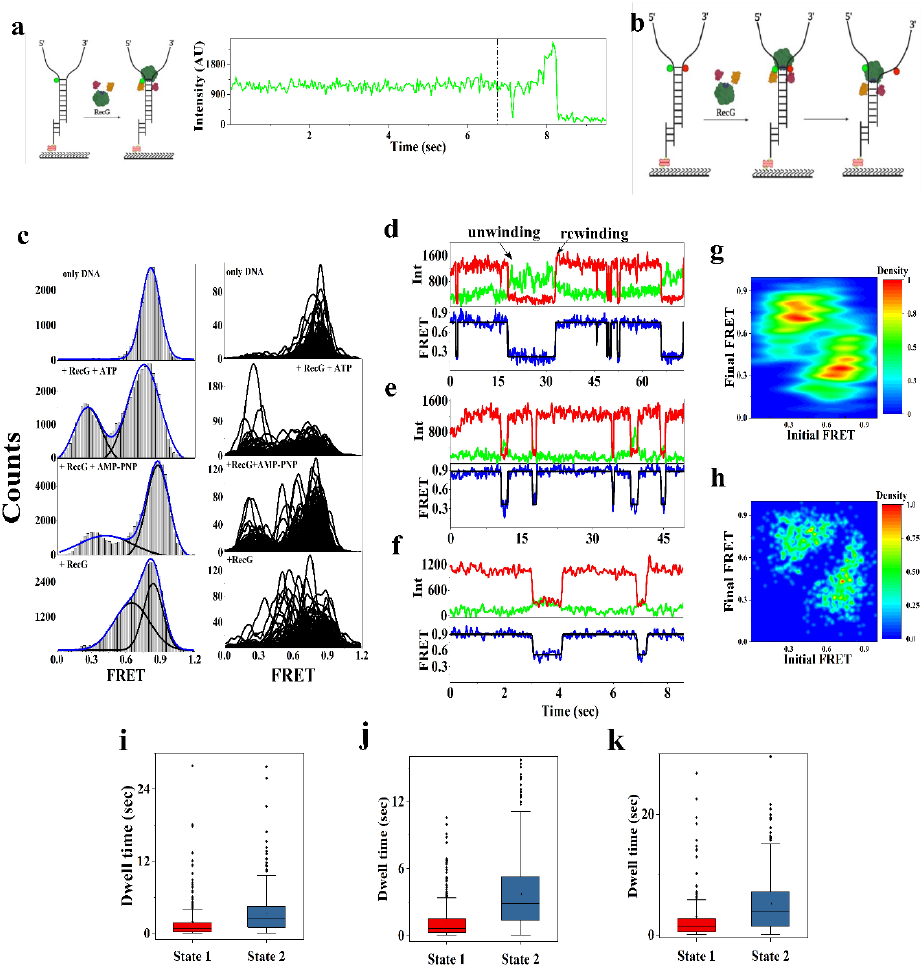
Reiterative unwinding and rewinding of open forked DNA structure by RecG **(a)** Schematic representation of smPIFE experiment where the junction is labelled with only donor (Cy3) and recording of a single molecule trace depicting enhancement of signal upon insertion of MtRecG (at point of dotted line). **(b)** Pictorial illustration of the unwinding of the ss/ds junction of fork DNA by RecG, where the junction is labelled with donor (Cy3) and acceptor (Cy5) at opposite strands. **(c)** FRET histograms of ensemble and individual molecules for only open forked DNA (first box), DNA bound with RecG and 5mM ATP (second box), 5mM AMP-PNP (third box) and without any supplements (fourth box). Representative smFRET traces demonstrating reversible zipping and unzipping events by RecG supplemented with **(d)** 5mM ATP **(e)** 5 mM AMP-PNP **(f)** without any supplements. Transition density plot (TDP) depicting dynamical transitions in the FRET states when the open forked DNA is treated with **(g)** RecG and ATP **(h)** RecG and AMP-PNP. Box plot of the dwell time distribution for the dynamic states of the DNA when allowed to bind with **(i)** RecG and ATP **(j)** RecG and AMP-PNP **(k)** RecG without any supplements. Here State 2 corresponds to the zipped DNA or high FRET state and State 1 is for the partially unzipped DNA or intermediate FRET state for (k), and fully unzipped DNA or low FRET state for (i) and (j). (Colour Coding: green for donor, red for acceptor, blue line depicts FRET Efficiency and Black line is to denote HMM fitted FRET state)

In order to extrapolate the dynamics of open fork DNA while bound to MtRecG, it is labelled with Cy3 and Cy5 placed opposite to each other in two different strands, exactly at the initiation point of the duplex DNA termed as junction and immobilization is carried out through extension of one of the duplex strands having complementarity with a biotin carrying DNA strand (Figure 1b). In previous studies it has already been verified that modified junction with fluorophores and the proximity of immobilization does not influence the activity of helicases.^44–46^ When MtRecG is not present in the system, there is predominancy of a single high FRET state (0.8) as inferred from the ensemble histogram (Figure 1c). However, there is occasional very short-lived (average dwell time being 0.43 ± 0.04 sec) switching to a low FRET state which varies from 0.2 to 0.3 FRET Efficiency in almost all the molecules as has been shown in the representative time trace and FRET histogram plot of the individual molecules (Figure 1c, Supplementary Figure S4). The occurrence of this low FRET state has been attributed to the phenomenon of molecular fraying, that occurs due to transient opening of the terminal base pairs of a double stranded DNA. Upon application of MtRecG into the sample chamber, we found noticeable changes in the FRET dynamics of the open fork DNA. Once MtRecG is flowed along with 5mM ATP, there were episodes of rapid decrease in the FRET state which remains unchanged for considerably longer amount of time followed by reappearance of the previous high FRET state as swiftly as it has decreased before (Figure 1d). The repeatedly varying FRET levels between two values can be seen in the single molecule time trace and the FRET Efficiency plot as shown in Figure 1d. This particular property of carrying out cycles of zipping and unzipping of base pairs in a reversible fashion by MtRecG is highly consistent with other helicases irrespective of the fact that whether they belong to the same superfamily or not.^47–49^ Through ensemble histogram two clear peaks, one at FRET state 0.26 and other at 0.75 are well reflected and those corresponds to the unzipped and rezipped state of the fork DNA respectively (Figure 1c). This oscillatory behavior of all the open fork DNA molecules as depicted in the graph portraying individual histograms is attributed to MtRecG catalyzed unwinding with subsequent rewinding. However, through the individual histogram plot we found exactly three exceptional molecules where the open forked junction has remained in the unwound state for considerably longer amount of time as compared to the zipped state (Figure 1c). The rewound state is characterized as the ‘waiting time’ for MtRecG since during that particular period the protein stalls and prepares itself for its next round of unwinding activity. The distribution of the dwell times at the two FRET states, where State 1 depicts the low FRET or ‘unwounded state’ and State 2 is for the high FRET or ‘rezipped state’, have been represented through box plot (Figure 1c). The average duration at the unwound state (State 1) is found to be τ_u_ = 1.84 ± 0.20 sec and characteristic waiting time (State 2) came out as τ_w_ = 3.38 ± 0.18 sec (Figure 1i). Besides, upon quantitative characterization of the switching between the two states through transition density plot, we found that maximum number transitions have taken place from a considerate low FRET state (0.3) to a high FRET (0.8) state or vice versa (Figure 1g). And, this observation denotes that there occurs complete local unwinding at the ss/ds DNA junction by MtRecG when supplemented with ATP. Next, to scrutinize the unwinding activity of MtRecG in more detail, the ATP in the system is replaced with the same concentration of AMP-PNP (non-hydrolysable analog of ATP) and we could still observe reiterative unwinding and rewinding of open forked DNA by MtRecG (Figure 1e). However, there is a shift in the E_FRET_ value towards higher FRET region (0.38) corresponding to the unwound state and the population also got diminished (Figure 1c). Besides, the dwell time at the unwound state also decreases to 1.24 ± 0.07 sec and alternatively, the rewound state displays a higher dwell time of 3.73 ± 0.15 sec (Figure 1j). The transition density plot also shows a shift in maximum transitions from high FRET state (0.7) to nearly intermediate state (0.48) and contrariwise. Further, in the absence of both ATP and AMP-PNP, the ensemble histogram exhibits complete disappearance of the low FRET state peak which depicts that without any external fuel, MtRecG is unable to perform the job of hydrogen bond breakage with full efficiency all by itself. And, other than the fully zipped high FRET state there is occurrence of an intermediate FRET state where the E_FRET_ value (0.65) is not substantially different from the fully rewound or high FRET state (0.81) (Figure 1c and 1f). The set of above-mentioned observations uncovers certain features of MtRecG about its helicase nature. (i) There is rapid unzipping of the junction base-pair by MtRecG, in presence of external fuel ATP and that is why we found an abrupt drop in the E_FRET_ value unlike the cases of other helicases where there is appearance of saw tooth pattern during the unwinding stage.^44,48^ However, the quick rewinding process that is again followed by unwinding after a pause coincides with the pattern executed by other helicases (ii) Even though the unwinding activity is dependent upon ATP, the exhibition of partially unzipped state in absence of ATP or any other ATP analog illustrates that there is residual unzipping activity of MtRecG in the absence of any energy source. (iii) The persistent re-initiation of DNA unwinding by MtRecG after reannealing denotes that the phenomenon of unwinding the full substrate is missing, rather after opening the bonds till a certain extent, MtRecG translocates to the original junction position with concomitant rezipping.

### Partial Unwinding of the junction of Fork DNA with a 3’ and 5’ single stranded overhang by MtRecG

The analysis of the helicase activity of MtRecG towards the stem region from the junction has been expanded further on partial forked DNA with a 3’ and 5’ overhang which act as intermediates of replication during occurrence of a lesion in one of the parental template strands. As mentioned in the previous section, the binding of MtRecG at the junction of a 3’ and 5’ overhang forked DNA is confirmed through smPIFE where the substrates, has been singly labelled at the junction with donor fluorophore (Figure 2a and Figure 3a). There is approximately two-fold increment in the fluorescent intensity of Cy3 signal within the very first second of flowing of MtRecG into the fluid channel (Figure 2a and 3a) which approves the attachment of protein. The labelling of the junction of fork with 3’ overhang, mimicking a structure of replication fork having only the nascent lagging strand, for smFRET has been carried out as explained in the previous section and represented in Figure 2b and Supplementary Figure S2b. In the absence of MtRecG, the fork with a 3’ single-stranded overhang demonstrates a single high FRET state (0.84) indicating the zipped state of the ss/ds DNA junction (Supplementary Figure S5 and Figure 2c). However, there is still a meagre population in the low FRET state as observed in open forked structure, confirming existence of molecular fraying at the double stranded termini even when one of the arms are blocked (Figure 2c). We found extensive change in the E_FRET_ profile of 3’ overhang forked DNA while being treated with MtRecG and 5mM ATP. There is appearance of two more FRET states, one having E_FRET_ value at an intermediate range, 0.66 and another at low FRET state (E_FRET_ = 0.3) in the ensemble histogram (Figure 2c). Besides, approximately all the molecules have resided in all the three dynamic conformations portrayed by high, intermediate and low FRET states as being imitated in the individual histogram plot (Figure 2d). The intermediate FRET state corresponds to ‘partially unzipped state’ of the fork and the low FRET state depicts ‘complete unwinding’. However, the integrity in the reiterative unwinding nature of MtRecG is conserved with a difference, that in case of a 3’ overhang forked DNA, there are frequent events of ‘partial unwinding’ accompanied by fewer episodes of ‘total unwinding’ along with ‘waiting period’ of MtRecG. The preference for partial unwinding is significantly reflected in the dwell time as well, where average existing time for the partially unzipped state (State 2) is τ_pu_ = 1.21 ± 0.08 sec which is almost two-fold higher than that of the fully unwound state (State 1), that is τ_u_ = 0.57 ± 0.06 sec (Figure 1i). Furthermore, the heat map of transition density plot also confirms that unwinding occurs heterogeneously by revealing roughly equivalent transitions from high to low, high to intermediate or intermediate to low FRET states and transitions happening in the other way round (Figure 2c). Next, when ATP in the system is replaced with AMP-PNP and when both ATP and AMP-PNP is removed from the system, we found no significant change in the E_FRET_ distribution of the ensemble or individual histograms, which demonstrates the same inclination towards ‘partial unwinding’ over ‘complete unwinding’ by MtRecG (Figure 2c, 2f and 2g). However, there is full consistency in continuous oscillation at non-periodic intervals of the FRET signals. Besides, the average dwell time in the ‘partially unwound’ state (State 2) also remains two-fold higher than that of the complete unwound state (State 1) in case of AMP-PNP supplement, which is 1.57 ± 0.09 sec versus 0.78 ± 0.07 sec (Figure 2j). The same trend follows when both the supplements (ATP and AMP-PNP) are removed from the system and the average dwell times come out as 1.07 ± 0.06 sec and 0.52 ± 0.06 sec for partially unzipped (State 2) and fully unzipped states (State 1) respectively (Figure 2k). In TDPs, either in presence of AMP-PNP or absence of any supplements we found unchanged transitions from all possible ways low to intermediate, intermediate to high and low to high FRET states with transition in the other way round as well (Figure 2h). Therefore, the results illustrate that the ss/ds DNA junction unwinding property of MtRecG for the replication fork with a nascent lagging strand is reiterative in nature being more persuaded towards ‘partial unzipping’ and is independent of any external fuel.

**Figure 2.**
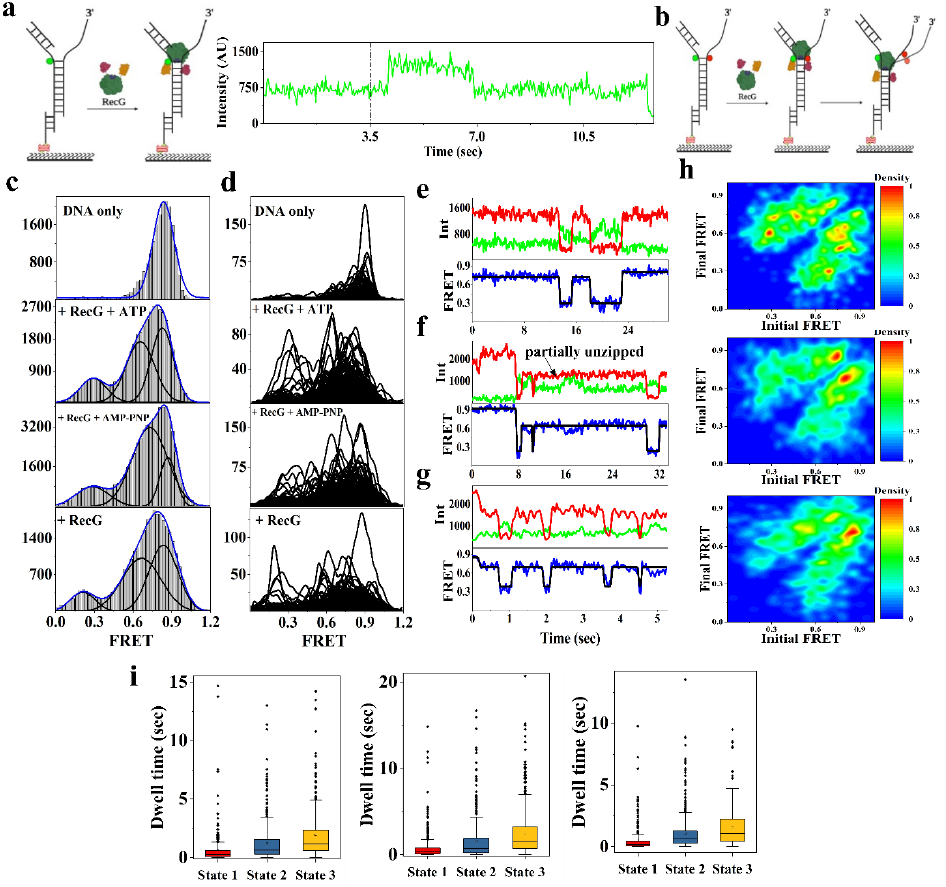
Partial unwinding of the junction of a 3’ overhang fork DNA by RecG **(a)** Pictorial representation of smPIFE experiment where the junction is labelled with donor and RecG is allowed to bind with the fork. Intensity versus time graph of the PIFE experiment, the dotted line denotes the point of insertion of RecG along with ATP. **(b)** Schematic model of partial unwinding of the junction of 3’ overhang forked DNA by RecG. **(c)** Ensemble FRET histograms of the 3’ overhang fork DNA when allowed to bind with RecG at different reaction conditions. **(d)** Overlapping FRET histograms of the individual molecules at different reaction conditions indicated in the box. Representative single molecule trajectory of the intensity of donor and acceptor versus time of 3’ overhang forked DNA when subjected to bind with **(e)** RecG and ATP **(f)** RecG and AMP-PNP **(g)** only RecG. **(h)** Transition density plots (TDP) displaying dynamical transitions of the 3’ overhang forked DNA when bound with RecG and ATP, RecG and AMP-PNP and only RecG. **(i)** Representation of the dwell time distribution at various states through box-plots where the first, second and third plot corresponds to the 3’ overhang fork bound with RecG and ATP, RecG and AMP-PNP and only RecG respectively. State 1 depicts the unwounded DNA or low FRET state, State 2 stands for partially unzipped state or intermediate FRET state and State 3 is for the fully wound or high FRET state.

**Figure 3.**
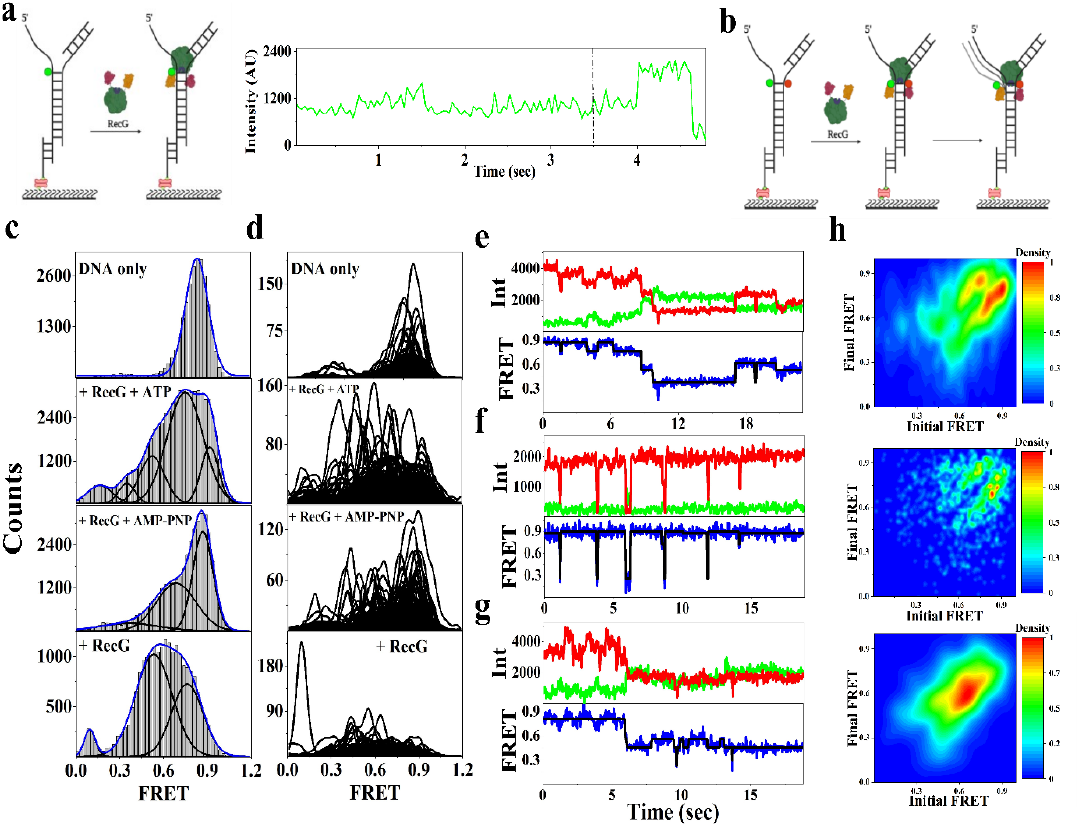
Inconsistent unwinding of the junction of a fork having a 5’ overhang. **(a)** Schematic diagram of smPIFE experiment and single molecule time trace of a molecule depicting increment in intensity of the Cy3 upon flowing the protein. The dotted line depicts the point of insertion of protein along with ATP **(b)** Illustration of the helicase activity of RecG on a fork DNA with 5’ overhang fork DNA. **(c)** Ensemble and **(d)** overlapping individual molecular FRET histograms depicting the FRET states of the substrate fork DNA with 5’-overhang, and when it is bound with RecG along with different supplements as mentioned in the box. Single molecule donor-acceptor intensity and FRET time trajectory of the substrate DNA when bound with **(e)** RecG and ATP **(f)** RecG and AMP-PNP **(g)** only RecG. **(h)** TDP depicting FRET transitions when the substrate is bound with RecG and ATP (top), RecG and AMP-PNP (middle) and only RecG (bottom).

The study of junction unwinding features have further been seen on the replication fork having a nascent leading strand consisting of forked DNA with a 5’ overhang through smFRET. The labelling scheme is represented in Figure 3b and Supplementary Figure S2c. Just like the previous two forked DNA substrate, here also we found occurrence of a single high FRET state (0.83) in the ensemble histogram (Figure 3c and Supplementary Figure S6) in the absence of protein. Then, delivery of MtRecG with and without various supplements brings changes in the dynamicity of the fork. In all the cases, along with the high and low FRET states there is appearance of multiple other partially unwound states reflected via the broad distribution in the ensemble histogram at intermediate E_FRET_ region (Figure 3c). There is no general trend observed in the change in dynamics of fork when the reaction buffer is supplemented with ATP or its non-hydrolysable analog or is rid of both. FRET histogram of individual molecules also portrays, that there is a huge population at the broad region of E_FRET_ = 0.4 - 0.7 with very few exhibiting the low FRET state (Figure 3d). Moreover, even there is presence of many states in the ensemble histogram we couldn’t find its reflection in TDP, especially for the molecules in the low FRET state were not found to go through any transition (Figure 3h). Now this observation points towards a possible conclusion that in some molecules there is non-reversible unwinding of the junction and that is independent of buffer system and we found disordered FRET histograms upon changing buffer supplements. Unwinding activity of any helicase is highly dependent upon the mode of translocation through its substrate. Thus, through our observations we concluded that translocation of MtRecG in 5’-3’ direction from the junction along the template leading strand of the replication fork is highly erratic in nature. And, that is why whenever the 5’ overhang forked DNA is subjected to bind with MtRecG irrespective of the fact that whether there is presence or absence of any supplements we found occurrence of some assorted intermediate FRET states corresponding to various random partially unzipped conformations. Furthermore, in previous studies, it has already been confirmed that the response of RecG is inhomogeneous with respect to the partial forks carrying either the leading or lagging nascent strands in terms of binding, translocation as well as helicase activity.^27,30^

### MtRecG does not exhibit its helicase activity at the stem region of Forked DNA

Next, we sought to investigate the extend of opening of duplex stem region of the fork since reiterative unwinding and winding events confirms no complete separation of the two strands by MtRecG. For this, the labelling position of the donor (Cy3) and acceptor (Cy5) is moved towards the stem region. The Cy3 is shifted to 3 nucleotides and Cy5 is shifted to 11 nucleotides away from the junction for forked structures having two single stranded overhangs and one 3’ overhang (Figure 4a and 4d). When the system is devoid of protein, the open forked DNA exhibited two dynamic, high and intermediate FRET states corresponding to 0.73 and 0.54 (Figure 4b). However, when MtRecG is flowed into the fluid chamber along with 5mM ATP we found no significant change in the E_FRET_ profile of the ensemble histogram, just having a broad distribution from intermediate to high FRET state regions (Figure 4c). The E_FRET_ histogram of the individual molecules also reinforces the fact that there is no unzipping towards the duplex region of the open fork (Figure 4c). Similarly, for the partial fork with a 3’ overhang, the FRET profiles of the ensemble histograms are almost equivalent irrespective of the fact that whether MtRecG is present or absent in the system (Figure 4e and 4f). Also, the FRET profiles of the individual molecules are also not significantly different (Figure 4e and 4f). This proves that the unwinding property of MtRecG from the junction towards the duplex stem component of the fork is distinctly very local in nature confined to initial one or two base pairs from the junction and that is why there is exhibition of a rapid drop in FRET Efficiency during unwinding by MtRecG when labelling is at the junction. In case of open forked structure since, we found multiple rounds of complete unzipping and rezipping events at the ss/ds DNA junction and that gives the inference that there lies a return point of MtRecG from the duplex region of the fork. And, through the results of experiments where the stem region is labelled, we can conclude that the return point is right from the second or third base pair from the junction. Furthermore, the waiting event of MtRecG characterized in the previous section during its open forked DNA’s junction unwinding action, now can be attributed to the phenomenon of ‘backward strolling’ of the motor protein along either of the single stranded overhangs. However, for a partial fork having a 3’overhang, we observed a higher tendency for partial unwinding followed by rewinding which confirms that right at the junction only MtRecG has lower unwinding processivity and hence the unwinding process is not taken forward towards the duplex stem unit. Thus, in case of partial fork with a 3’-overhang as well we are able to conclude that during the waiting period of junction unwinding event, specifically ‘retrograde motion’ of MtRecG was being exhibited.

**Figure 4.**
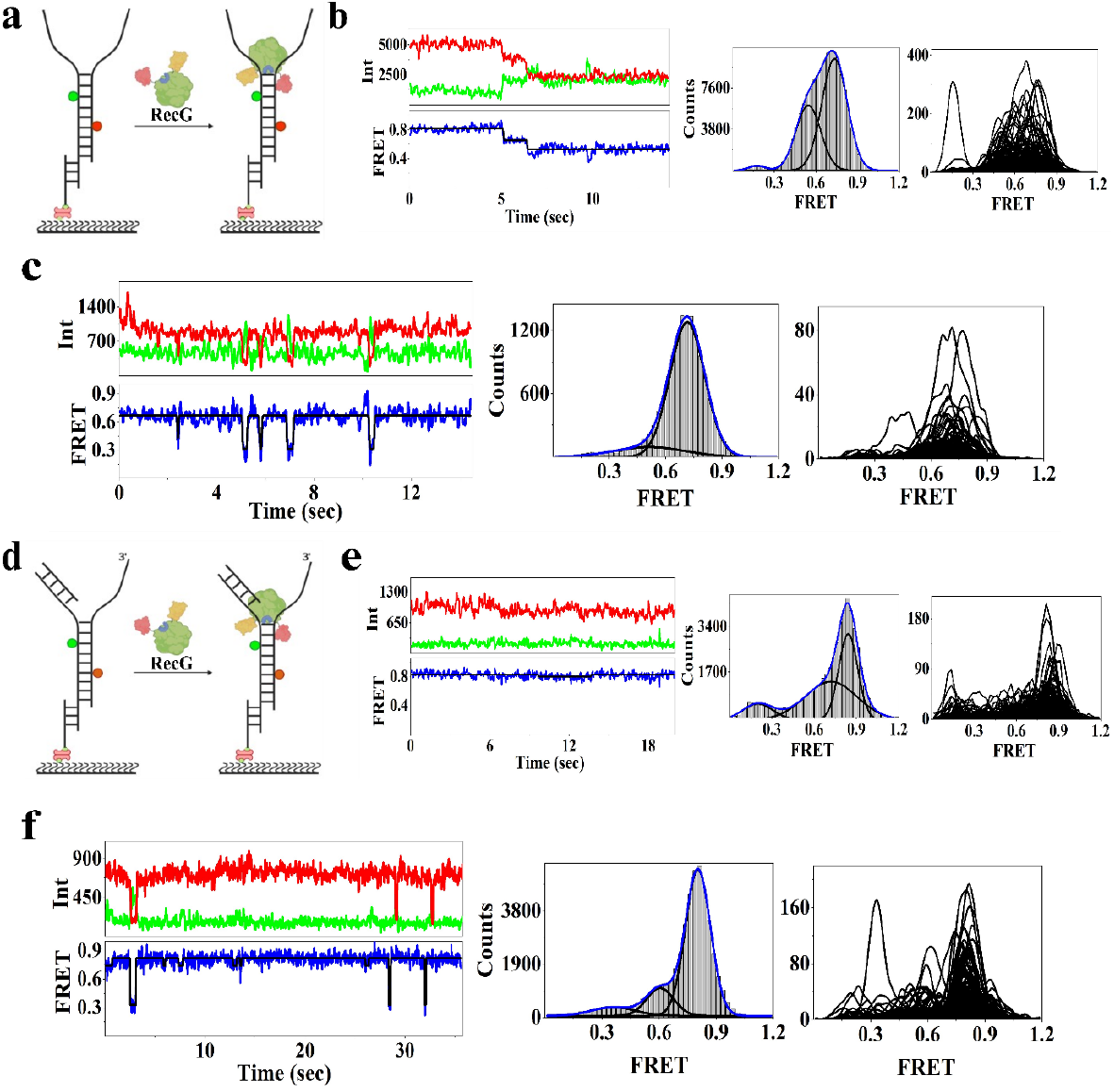
Non occurrence of unwinding action of RecG towards the stem region of the fork. Schematic illustration of smFRET assay for depicting the helicase action of RecG at the duplex stem region of the **(a)** open forked DNA and **(d)** forked DNA with a 3’overhang. smFRET and FRET Efficiency time trace, ensemble and individual molecular FRET Efficiency histogram of **(b)** open forked DNA labelled at the stem region **(c)** open forked DNA labelled at the stem region subjected to bind with RecG **(e)** fork DNA with a 3’ overhang labelled at the stem region **(f)** fork DNA with a 3’-overhang labelled at the stem region allowed to bind with RecG

### Fork Reversal Activity of MtRecG is more processive on the partial fork substrate having a 3’ overhang

Through several previous studies it is known that MtRecG plays a key role in modification of a stalled replication fork that can eventually facilitate replication restart. And, the unique role of MtRecG lies on the point that it has the capacity to promote fork reversal by unwinding the newly replicated arms of the fork from their corresponding parent strands. However, the direct experimental information in terms of quantity about the mode of local perturbation of DNA, helicase activity and translocation of the protein over forks carrying one of the nascent strands is missing. Through the results of the previous section, we got the signature of backward movement of MtRecG. Now, to specifically investigate the fork reversal event at each of the nascent strands separately, firstly the forked substrate is generated with a 3’ overhang by blocking the 5’ overhang region with its complementary oligonucleotide, arm2 (Supplementary Table 1). The designed substrate resembles a stalled fork where lagging strand synthesis has moved ahead than that of the leading strand. The donor is labelled at the junction of the template strand and acceptor is placed at the end of arm1 facing the ss/ds junction (Figure 5a and Supplementary Figure S2d). To mimic the crystal structure DNA substrate, a one base pair gap was left between the daughter lagging strand and junction of the fork (Supplementary Table 1). In the absence of protein, only DNA exhibits three peaks having E_FRET_ values of 0.48, 0.62 and 0.74 depicting high intrinsic fluctuation of arm2 (Figure 5b and Supplementary Figure S7). However, the rich dynamics observed between the template and the complementary arm2 strand is not critical for the purpose of this study. Hence, we moved forward by allowing the substrate DNA to interact with MtRecG supplemented with ATP. Once MtRecG is inserted in the chamber along with ATP, there is reappearance of alternative zipping and unzipping phenomenon and the zipped state holds the intrinsic fluctuation of the nascent strand (Figure 5c). The ensemble E_FRET_ profile shows a fresh peak at the low FRET region (0.21) along with three more peaks, of values 0.38, 0.61 and 0.77 (Figure 5c) which are almost similar to the FRET states of only DNA substrate. The series of observations indicates moving away of the protein from the labelled region since the intrinsic fluctuation of arm2 is getting resumed in the rewinding stage. Also, reiterative unwinding again, confirms that there is no complete separation of arm2 strand from its corresponding template DNA. Now, when MtRecG is allowed to act on the DNA without ATP, we found disappearance of the rises and dips of FRET states in smFRET traces, rather there are events of prolonged stabilized low FRET states with small interruptions of intermediate FRET value (Figure 5b and 5d). There is also a stark difference in the average dwell time of the unzipped state (State 1) in presence and absence of ATP, turning out to be 1.00 ± 0.19 sec (Figure 5g) versus 9.92 ± 1.08 sec (Figure 5h) respectively. Prolonged low FRET state proves longer unzipped state and intermediate FRET state denotes subtle movement of MtRecG promoting partial zipping. Further, both the ensemble and set of individual E_FRET_ histograms denotes peak position at two FRET states with E_FRET_ = 0.15 and 0.39 (Figure 5b). Therefore, this observation gives us the direct conclusion that the motion of MtRecG is getting stalled at the interface of the arm and stem region of the fork when the external fuel, ATP is removed from the system. Thus, direct inference from our results is, energy derived from ATP is utilized for mobility of MtRecG from the junction along the arm of the fork having the hybrid duplex of template and nascent lagging strand, since movement through the stem region of the fork has already been ruled out in previous section.

**Figure 5.**
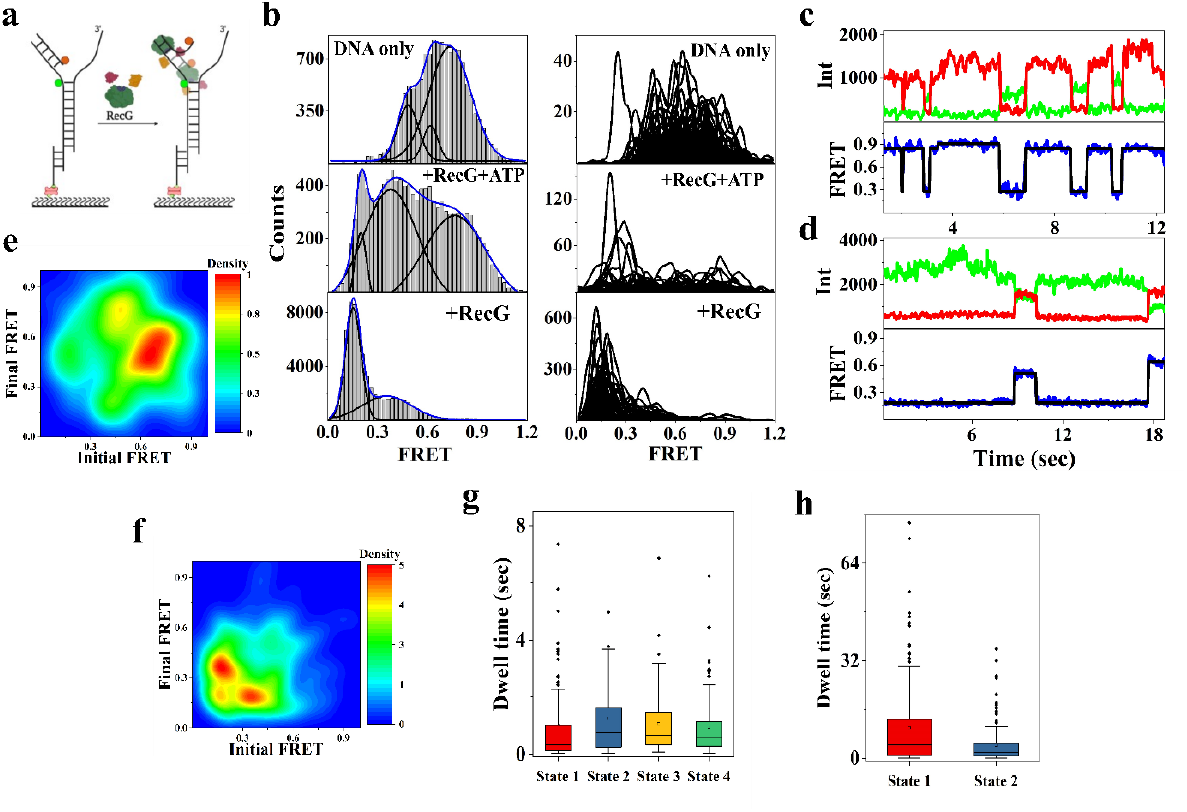
Reiterative fork reversal activity of RecG over the nascent lagging strand **(a)** Schematic diagram of the Cy3 (at the junction, green) and Cy5 (at the nascent strand terminal facing the junction, red) labelled fork DNA to assess the fork reversal property of RecG. **(b)** FRET histograms depicting ensemble and individual molecules for the fork DNA alone, when the fork is allowed to bind with RecG supplemented withs 5mM ATP and when the fork is allowed to bind with only RecG. Representative time traces of fluorescence emission intensities of donor and acceptor and FRET trace of the fork DNA when bound with **(c)** RecG and ATP **(d)** only RecG. Heat Map showing FRET transitions of the fork DNA when bound with **(e)** RecG and ATP **(f)** only RecG. **(g)** Box plot representation of the dwell time distribution at four different FRET states, when the fork is bound with RecG and ATP. State 1 stands for fully unzipped state (E_FRET_ = 0.2), State 2 for the intermediate state being more inclined towards unzipped form (E_FRET_ = 0.38), State 3 is for the intermediate state, more inclined towards the zipped form (0.59) and State 4 stands for the fully zipped state (0.8). **(h)** Boxplot of the dwell time distribution at the two FRET states when only RecG is bound with the fork DNA. State 1 stands for the fully unwound state and State 2 is for the intermediate state being primarily unzipped.

Now, the degree of fork reversal by MtRecG has been investigated on a fork with a 5’ overhang and the 3’ overhang region is blocked with arm1 (Supplementary Table 1 and Figure 6a) and the structure so formed resembles a fork where there is presence of only leading strand. The mode of labelling has been described in Supplementary Figure S2e. Here we found, that MtRecG is able to partially unwind arm1 from it’s cognate template strand since the peak positions of FRET histogram appeared at intermediate states (0.54 and 0.71) along with a high FRET state in presence of protein and ATP (Figure 6b and 6c) in contrast to only high FRET state when no MtRecG is there in the system (Figure 6b). Also, through TDP we got that there is primary transition from E_FRET_ value of 0.74 to E_FRET_ value of 0.51 and vice versa (Figure 6f). Further, when ATP is removed from the system the major Gaussian peak is at high FRET state with two minor peaks in the intermediate region (0.55 and 0.74) (Figure 6b) which are same as has been observed in presence of ATP however there is a decrement in population (Figure 6b and 6c) and major transitions are also in the region of high FRET (E_FRET_ = 0.89) to intermediate FRET (E_FRET_ = 0.75) and in the other way round (Figure 6g). Even though there is a shift in population towards lower FRET state in the ensemble histogram once MtRecG is bound with DNA, the population at the low FRET state which corresponds to complete unzipping, is significantly low indicating low processivity of MtRecG while unwinding the arm1 from its complementary unit. And these observations depict that the mode of erratic translocation of the helicase remains conserved for the template DNA having 5’-3’ directionality from the junction, even during reverse track motion. Thus, through our results we could decipher that MtRecG exhibits a clear asymmetric functionality towards fork reversal for leading and lagging strand. And, these signatures suggest that MtRecG faces both translocation based as well as processivity issue while locomoting along 5’-3’ directed template strand from the junction. So, we can conclude that formation of ‘chicken foot intermediate’ through fork regression by MtRecG does not occur via continuous and symmetric unwinding and rewinding of both the nascent daughter strands. Rather, we found that the fork reversal activity of MtRecG is prominently one sided for the kind of stalled forks where the lagging strand synthesis has continued beyond the leading strand and there lies a gap in the leading strand region consistent with several other ensemble studies and the crystal structure.

**Figure 6.**
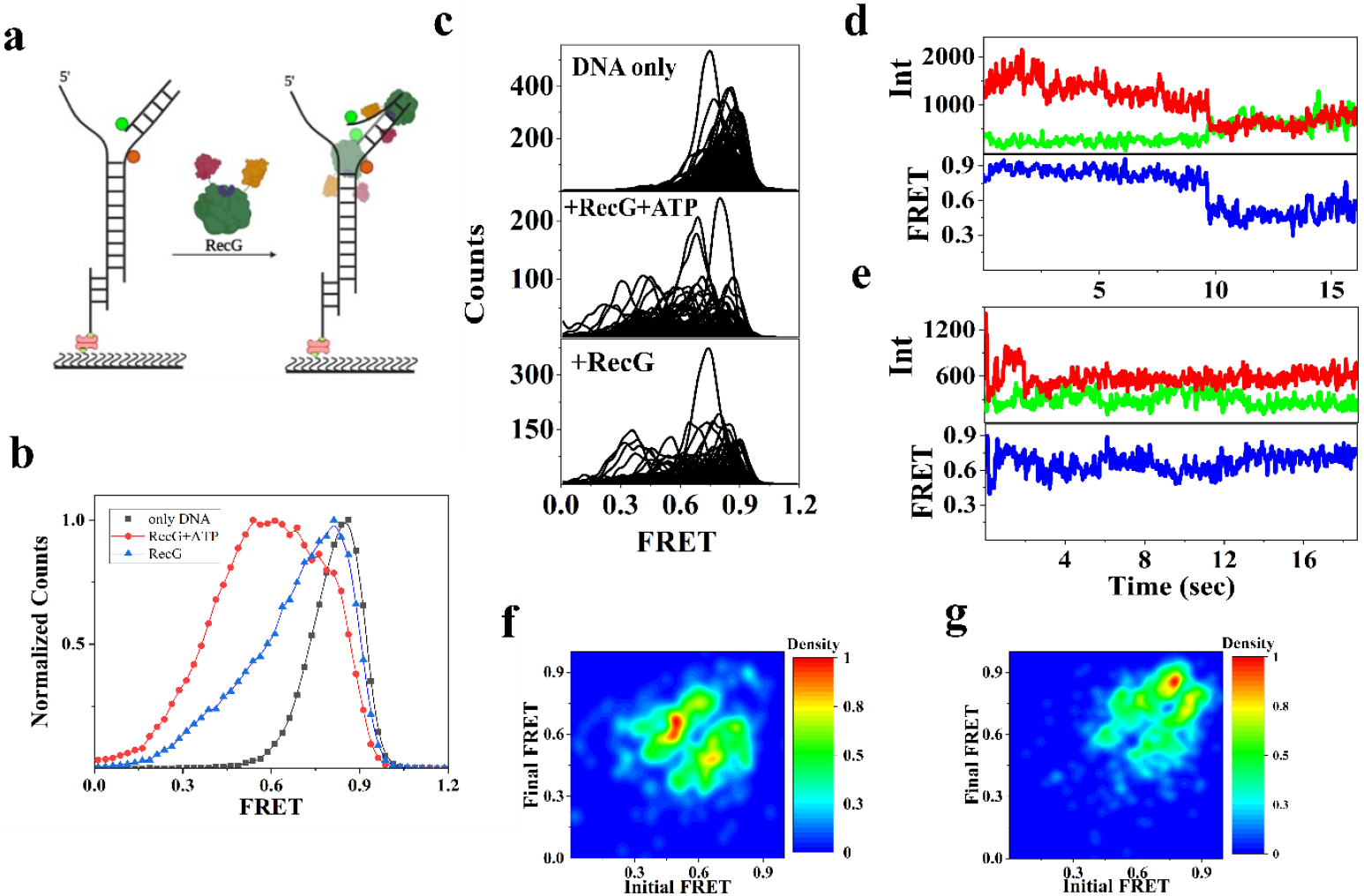
Low processivity of RecG in unwinding the leading daughter strand from it’s template DNA **(a)** Schematic of the smFRET assay exhibited for verifying fork reversal along the leading strand by RecG. **(b)** Ensemble FRET histograms of the substrate DNA, DNA with RecG and ATP and with only RecG. **(c)** Overlapping FRET histograms of the individual molecules with reaction conditions mentioned in the box. Donor and acceptor intensity, and FRET Efficiency vs time trace of the DNA substrate when allowed to bind with **(d)** RecG and ATP **(e)** only RecG. Heat Map demonstrating FRET transitions of the substrate when bound with **(f)** RecG and ATP **(g)** only RecG

### Non reversible Unwinding of arm 2 in a blocked fork structure by MtRecG

A heterologous stalled replication fork has been designed where both the daughter strands has come to halt simultaneously. However, the junction designed for experimental purpose is non homologous in nature to circumvent the possibility of non-specific annealing which can be a hindrance for a stabilized structure. However, due to non-complementarity in the arm region tracking the phenomenon of reannealing of the unwound daughter strands is not conceivable. The binding of MtRecG at the junction of a blocked fork DNA is verified through smPIFE (Figure 7a and Supplementary Figure S3) and we found that there is spike in the intensity ensuring clamping of the protein close to labeled fluorophore. The labeling scheme for monitoring the fork reversal activity of MtRecG on a three-way duplex forked structure has been described in Figure 7a and Supplementary Figure S3f. During unwinding of arm 2 in a blocked replication fork we found very distinctly that there is functional switching of MtRecG from ‘reiterative mode’ to ‘highly processive mode’ exhibiting irreversible unzipping of the nascent lagging strand characterized through occurrence of low FRET state (∼0.2) (Figure 7d and 7e). To verify the procedure quantitatively, we performed time-dependent assay taking movies at an interval of 2 minutes at different fields of view in the same fluid channel having the DNA substrate with MtRecG and ATP (Figure 7c). The consistency in reiterative FRET changes is broken and there is progressive disappearance of Cy5 spots with time indicating permanently unzipped DNA strand. As a control, in the blocked fork DNA substrate where MtRecG is not added, we found no significant FRET changes and minimal decrement in the Cy5 spots confirming that the disappearance of spots in the Cy5 channel is due to strong activity of MtRecG on a blocked fork (Figure 7f). Several long movies were also recorded and through deep analyzation of individual molecules it has been revealed that there is a rapid fall in E_FRET_ value when the substrate is subjected to the bind with protein. Prolonged low FRET state confirms that arm 2 is unzipped irreversibly from the template and it is in a stable single-stranded form for making itself available towards further modification. Nevertheless, in few molecules we found reappearance of higher FRET state after a long period of time, and that portrays either the protein is getting dislodged (Supplementary Figure S9) or there is reannealing of the template and arm 2 hybrid strands due to unavailability of any scope for finer molecular amendments. However, average dwell time is found to be 13.13 ± 1.19 sec, turning out to be higher than that of the unwound state in other cases and hence it is indicated that there can’t be further rewinding rather the protein is falling off. Next, the unwinding activity of MtRecG has been assessed for arm 1 as well which is a part of blocked forked DNA (Figure 7g and Supplementary Figure S2g) and we found no significant change neither in the number of Cy5 spots nor in FRET profile with time (Figure 7g).

**Figure 7.**
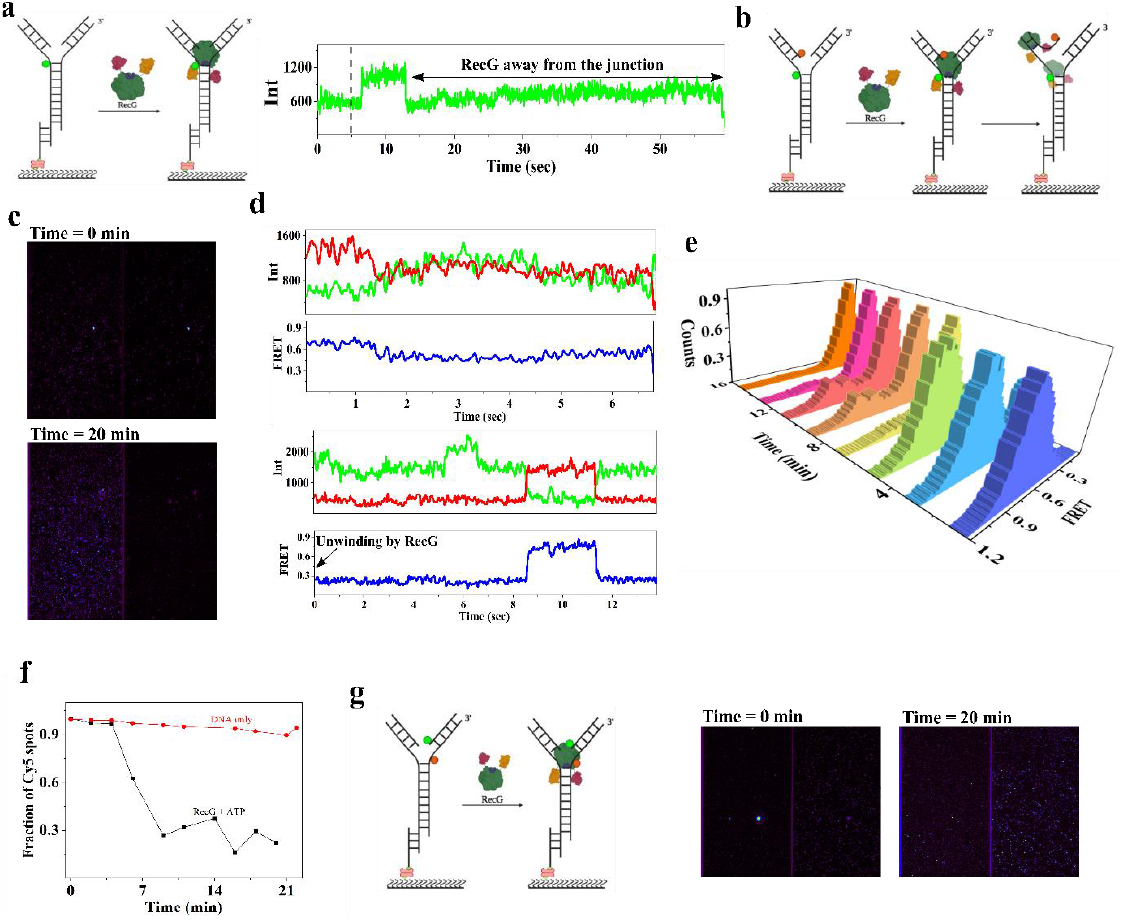
**(a)** Schematic diagram representing the set-up of smPIFE assay for the blocked fork structure, single molecule time trace of the smPIFE depicting enhancement in intensity once the protein is flown into the system. The intensity resumes to the initial level after a certain amount of time depicting movement of RecG away from the junction. **(b)** Schematic illustration of smFRET study for blocked fork DNA when allowed to bind with RecG **(c)** Lagging strand unwinding as monitored by Cy5 spots, when RecG is just flown into the system and 20 minutes after the introduction of RecG. **(d)** Single molecule donor and acceptor (Cy3 and Cy5) intensity and FRET Efficiency versus time trace of individual representative molecule at time = 0 min (first trace) and at time = 20 min (second trace). **(e)** 3D representation of the FRET Efficiency histogram, where z-axis denotes time. **(f)** Plot of fraction of remaining Cy5 spots as a function of time for only blocked-fork DNA (red) and when it is allowed to bind with RecG + ATP (black). **(g)** Schematic illustration of smFRET study of nascent leading strand unwinding by RecG. Image of a section of the sample at time = 0 min and at time = 20 min.

This observation depicts that even after being exposed to both the daughter strands simultaneously MtRecG maintains its integrity towards distinct preference for lagging strand unwinding. Hence, we conclude that during fork regression by MtRecG there is no symmetrical or simultaneous unfastening and consequential reannealing of both the daughter strands so as to drive the stalled replication fork towards a Holliday junction conformation. Instead, there is only one-way unzipping of the daughter lagging strand which brings into consideration the obvious query, that how the stalled fork gets modified in the form of Holliday junction making itself a suitable target for RuvC.^13^ There lies the possibility of spontaneous branch migration as has been proposed earlier, once one of the nascent strands gets unwounded and if there is junction homology, there occurs unprompted unfastening of the second daughter strand and eventually, they both get annealed to facilitate a stalled fork towards Holliday junction formation.^27^

## Discussion

The nature of the structure of a substrate DNA is highly responsible for the functional response of any protein and we found that MtRecG is not an exception. Depending upon the kind of DNA lesion or damage checkpoint, the morphological features of a stalled replication fork is likely to get modified and MtRecG is found to manipulate its functional activity accordingly. With a break in one of the template strands, the mobility of the corresponding daughter strand gets affected while the movement of the opposite nascent strand is unperturbed. In those cases, we are supposed to get a replication fork arrested in form of having either of the 3’ or 5’ overhangs. Similarly, there can be cases where both the daughter strands have halted at the same time giving rise to a blocked fork substrate and likewise junctions where progression of both the strands have come to a halt at a distance, there we get forked structure with two overhangs. In this study, we have tried to unravel the underlying mechanism of aberrant helicase activity of MtRecG in all kinds of synthetic non homologous stalled fork structures resembling intermediates of replication, repair and recombination. And, functional characterization has been exemplified through quantifying the response of the substrate in terms of their molecular heterogeneity and dynamical response. Acting as an atypical helicase, MtRecG has been found to meticulously utilize the properties of a helicase for promotion of the conformation of Holliday junction. In every kind of stalled forks, we found that the recruitment of MtRecG is taking place at the junction and through initial clamping at the junction the motor protein exhibits its translocation coupled with the function of unwinding and rewinding. For a fork with two single-stranded overhangs, we found short-ranged but efficient classical molecular zipper-like helicase activity (Figure 8a) of MtRecG at the junction towards the stem region, executing unwinding and reannealing cycles in an ATP dependent manner. Furthermore, to our knowledge for the first time we deduced that this reverberating unwinding process is very local in nature limited to 2-3 base pairs from the junction towards the stem region as compared to 25 base pairs in case of HIM-6, belonging to RecQ family. And, this short-distance unwinding is also manifested through a rapid drop in FRET efficiency profile just like in the case of eIF4I/eIF4H complex proteins.^50^ Thus, even when there is non-availability of double stranded reverse track, the translocation of this particular helicase is confined to a very local region in the stem part of the fork. Now, combining our results we can deduce that there is a patrolling motion of MtRecG along the 5’-overhang in search of a suitable target to exhibit its helicase action. However, once there is introduction of a nascent lagging strand, the same protein exhibited a dissimilar unwinding process exactly at the junction giving more preference to a partially unwound state. By precisely sensing the available reverse track along the template-daughter duplex region, MtRecG helicase changes its mode of functional dynamics and allows itself to exhibit retrograde translocation. Our data revealed that before sliding back along the parent-daughter hybrid track it is important for MtRecG to relax the base pairs of the ss/ds junction. There is adoption of the mechanism of ‘active translocation and passive unwinding’ during its ‘sliding back’ mode along the lagging daughter strand since, in the absence of ATP we found that MtRecG is able to carry out the unwinding process quite efficiently however the occurrence of rezipping events become less frequent. Now, this observation provides the clue about why MtRecG has been selected to be an ATP-dependent helicase, as rezipping of a non-modulated ssDNA is necessary to protect it from various external damages. The sliding back mode of MtRecG, is slanted along the direction of lagging daughter strand and alternatively when there is presence of only the leading strand, MtRecG faces processivity issue and unwinding in the reverse direction is not efficient even in presence of external fuel (ATP), promoting predominance of partially unzipped daughter leading strand. During a lesion in the leading template strand, the synchronization between the two DNA polymerases gets uncoupled and the lagging daughter strand moves ahead leaving a gap in the leading strand region and these kinds of stalled forks act as a potential substrate for binding of MtRecG. In such types of forks, as has been hypothesized before, that the fork reversal activity of MtRecG promotes ‘template switching’ for nascent leading strand causing the unwounded lagging strand to act as template for continuation of leading strand synthesis (Figure 8b).^25,27^ Now, the newly synthesized leading strand is zipped with the lagging strand generating a four-way duplex structure which eventually rearranges itself to take the form of ‘chicken foot intermediate’ followed by Holliday junction. Here, we found that unlike other helicases for example, BLM and Pif1, where the helicases reel backwards to resolve a DNA secondary structure^47,49,51^, the backward motion of MtRecG is to propel the DNA secondary structure formation, Holliday junction. Thus, we characterize MtRecG as ‘secondary structure regenerating helicase’. Further, in the kind of forks where replication arrestation due to double strand breaks has caused synthesis of both the infant strands to get paused simultaneously, we found even being exposed to a symmetrical substrate catalytic unwinding is prompted unidirectionally along the lagging strand. It is widely known that in general, the mode of migration of the helicase through the template strand is coupled with peeling out of the complementary daughter strand. Hence, by corollary we can propose that the gliding of MtRecG from the junction through the two template strands is unequivocal and asymmetrical in nature being more prevalent along the template with 3’-5’ polarity from the junction. In previous studies through simulation, it has already been verified that in case of superfamily (SF) 1 and 2 DNA helicases asynchronous unwinding strongly corroborates with higher efficiency.^52^ Next, the hypothesis comes forward, when the lagging daughter strand is in unzipped state through spontaneous branch migration the leading strand drives itself to bind with the single stranded lagging strand (Figure 8c) and eventually the structure of Holliday junction is achieved facilitating a recombinational repair specific intermediate for the action of RuvC protein.^27^ However, vital insights into the exact dynamical details of rearrangement of hydrogen bonds to promote Holliday junction is not clear. Our results offer motivation for further studies on the next step of recombinational repair which promotes collapse of the stalled fork, making it suitable to continue the process replication.

**Figure 8.**
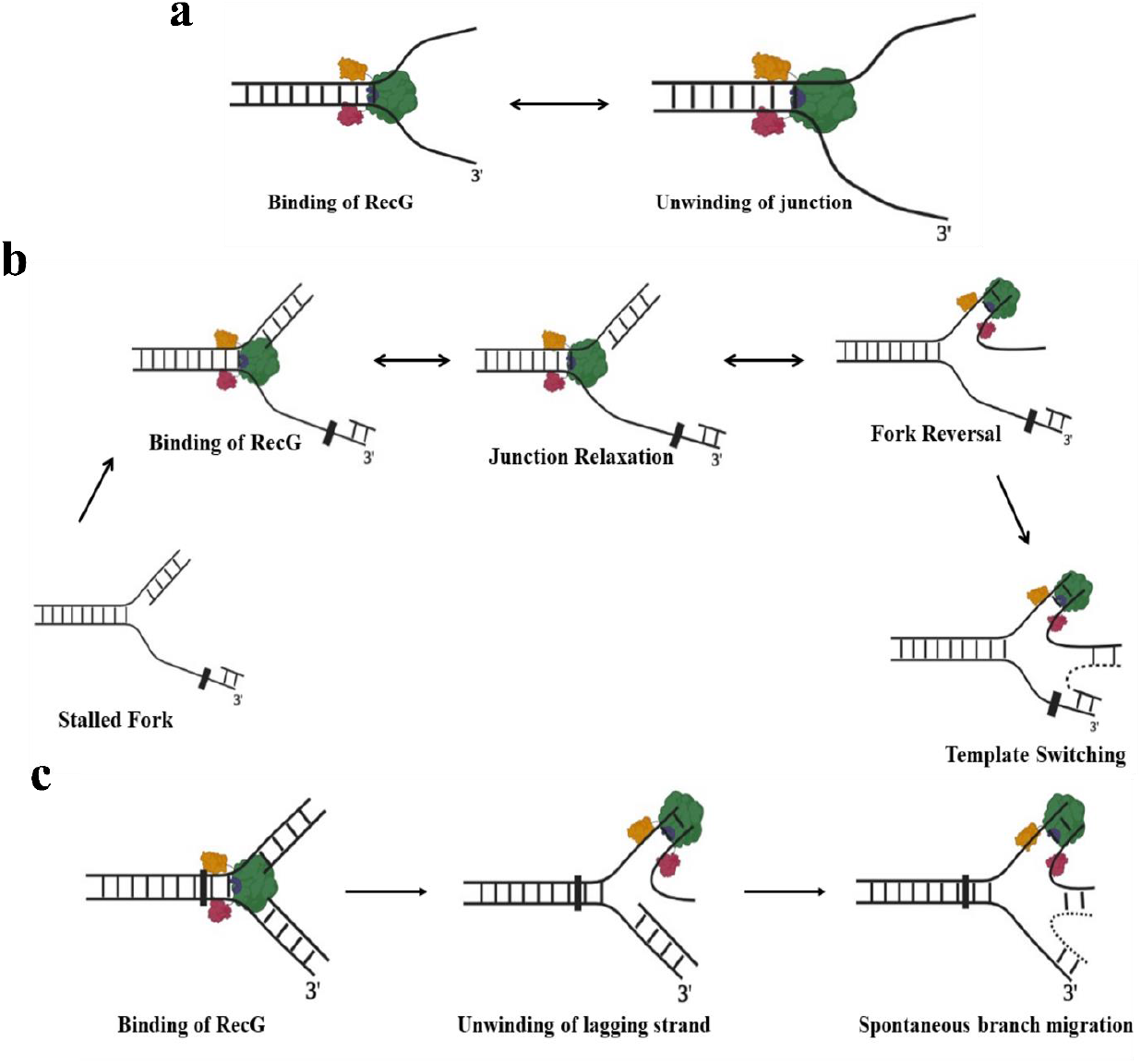
Asymmetric Fork Reversal Activity by RecG **(a)** Classical Molecular Zipper like model of RecG **(b)** Binding of RecG at the junction where there is a gap at leading strand region and unidirectional fork reversal **(c)** Asymmetrical unwinding of the lagging strand by RecG in a blocked fork

## Acknowledgements

The authors want to thank Soumen Mondal and Saikat Sadhukhan for helping with the writing of Matlab Codes, Debayan Purkait and Tapas Paul for their critical comments on the manuscript. This work is supported by DAE and DST-SERB funding agencies.

