## Supplementary Information for "Single Molecule Analysis of differential functional mechanisms of MtRecG while being exposed to variants of stalled replication-fork"

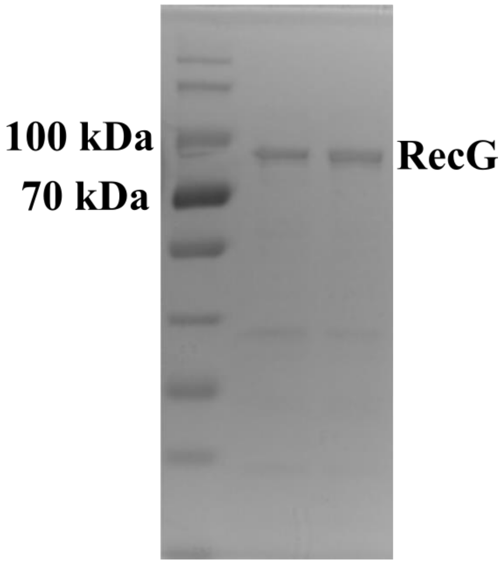


**Figure S1**. SDS PAGE Gel picture of purified RecG. We found band corresponding to 80 kDa molecular weight.

**Supplementary Table 1. DNA Oligonucleotides used for smPIFE and smFRET experiments**

| Construct Name | Sequence (5'-3') |
|---|---|
| Biotin-DNA | **BIOTIN**-AAAAATGTGTGTGTG |
| Strand1/ForkCy3 | GGCCAAAAAAG/**Cy3**/CATTGCTTATCAATTTGTTGCACCACACACACAC |
| Strand2/ForkCy5 | GGTGCAACAAATTGATAAGCAATG/**Cy5**/CGGCGCGATAT |
| Strand1/Fork | GGCCAAAAAAGCATTGCTTATCAATTTGTTGCACCACACACACAC |
| Strand2/Fork | GGTGCAACAAATTGATAAGCAATGCGGCGCGATAT |
| Arm1 | ATATCGCGCC |
| Arm2 | TTTTTGGCC |
| Arm2Cy5 | **Cy5**/TTTTTGGCC |
| Arm1Cy3 | ATATCGCGCC/**Cy3** |
| Strand3/ForkCy3 | GGCCAAAAAAGCAT/**Cy3**/TGCTTATCAATTTGTTGCACCACACACACAC |
| Strand4/ForkCy5 | GGTGCAACAAATTG/**Cy5**/ATAAGCAATGCGGCGCGATAT |

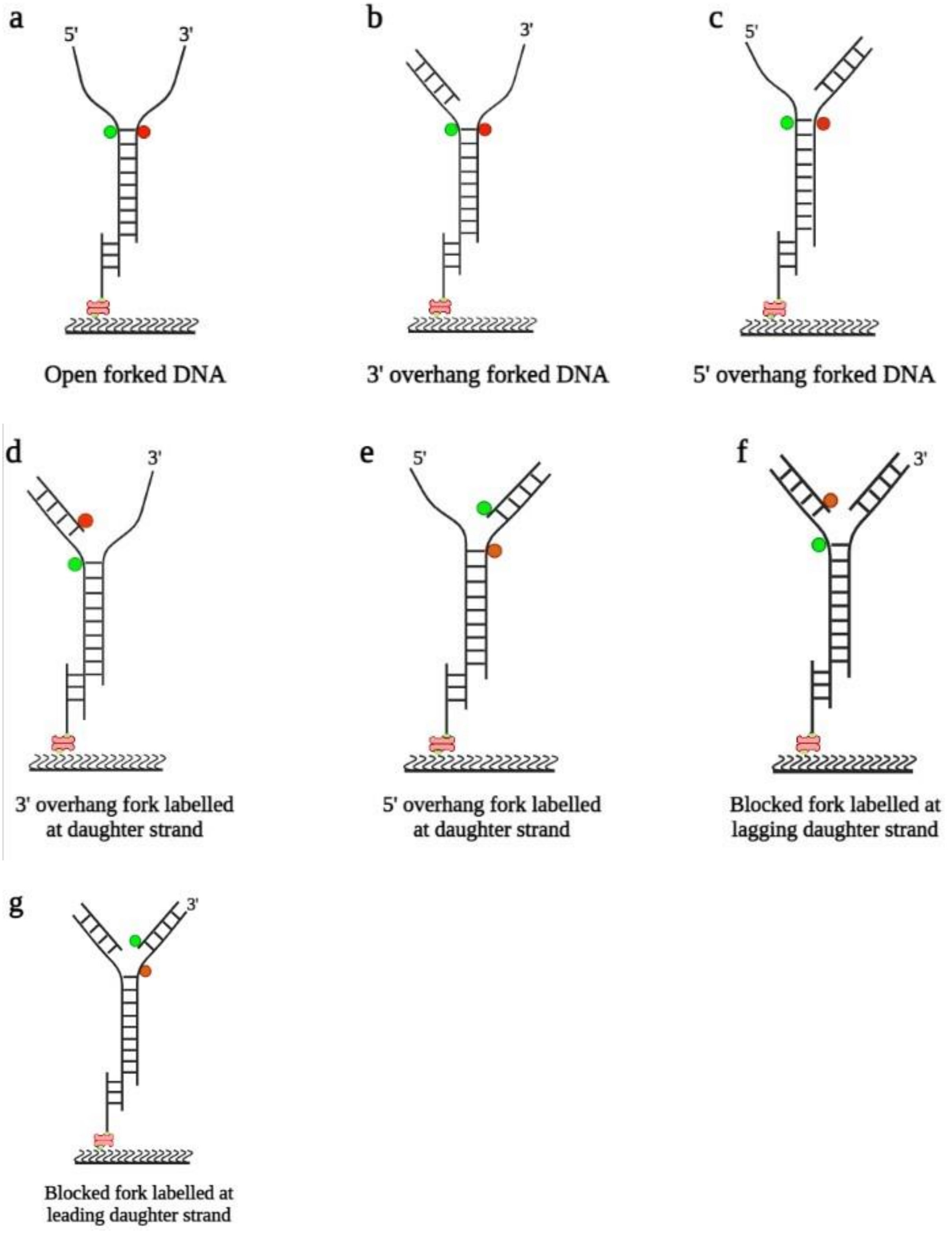


**Figure S2**. Labelling Scheme of the different DNA constructs used for smFRET experiments.

The DNA constructs used for smFRET experiments has been named as follows:

1. **Open forked DNA labelled at the junction**: Strand1/ForkCy3, Strand2/ForkCy5 and Biotin-DNA are annealed together to generate open forked DNA.
2. **3'-overhang forked DNA labelled at the junction**: Strand1/ForkCy3, Strand2/ForkCy5, Arm2 and Biotin-DNA are annealed together to generate 3'-overhang forked DNA.
3. **5'-overhang forked DNA labelled at the junction**: Strand1/ForkCy3, Strand2/ForkCy5, Arm1 and Biotin-DNA are annealed together to generate 5'-overhang forked DNA.
4. **3'-overhang fork DNA with acceptor labelled at the daughter strand**: Strand1/ForkCy3, Strand2/Fork, Arm2Cy5 and Biotin-DNA are annealed together.
5. **5'-overhang fork DNA with donor labelled at the daughter strand**: Strand1/Fork, Strand2/ForkCy5, Arm1Cy3 and Biotin-DNA are annealed together.
6. **Blocked fork labelled at lagging daughter strand**: Strand1/ForkCy3, Strand2/Fork, Arm2Cy5, Arm1 and Biotin-DNA are annealed together.
7. **Blocked fork labelled at leading daughter strand**: Strand1/Fork, Strand2/ForkCy5, Arm1Cy3, Arm2 and Biotin-DNA are annealed together.
8. **Open forked DNA labelled at the stem region**: Strand3/ForkCy3, Strand4/ ForkCy5 and Biotin-DNA are annealed.
9. **Fork DNA with a 3'-overhang labelled at the stem region**: Strand3/ForkCy3, Strand4/ForkCy5, Arm2 and Biotin-DNA are annealed.

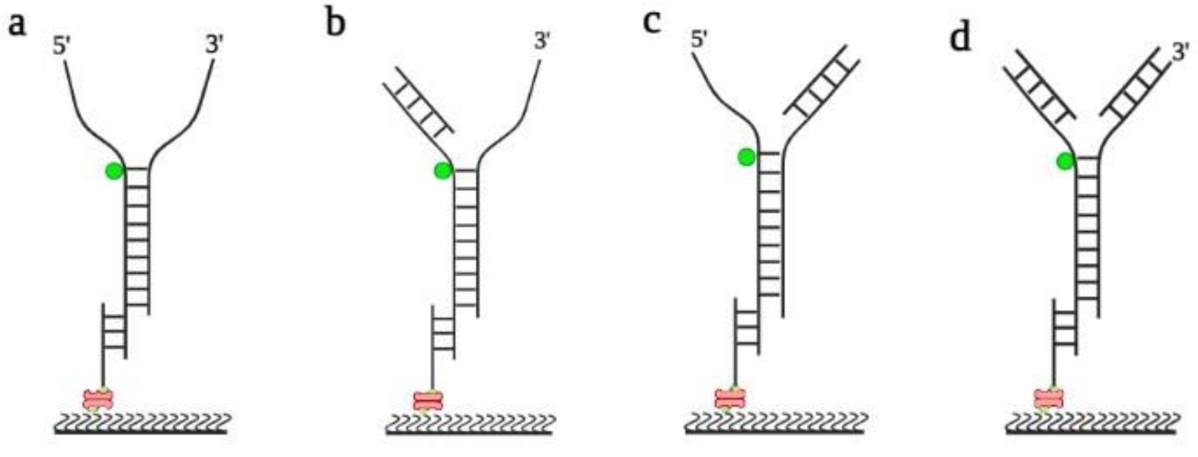


**Figure S3**. Labelling Scheme of the DNA constructs utilized for smPIFE experiment

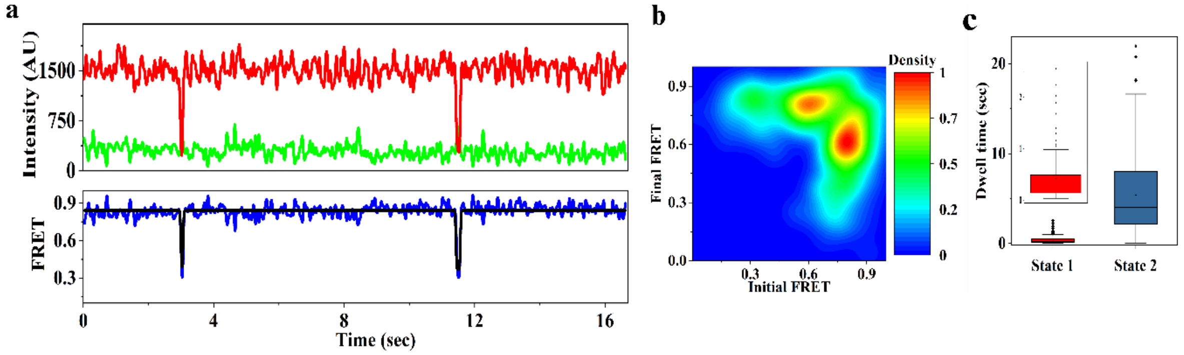


**Figure S4** (a) Representative single molecule time trace of the donor and acceptor intensity and FRET Efficiency versus time of open forked DNA labelled with donor and acceptor at the ss/ds junction (b) Transition density plot and (c) Box plot of the dwell time distribution, State 1 depicts low FRET state and State 2 depicts high FRET state

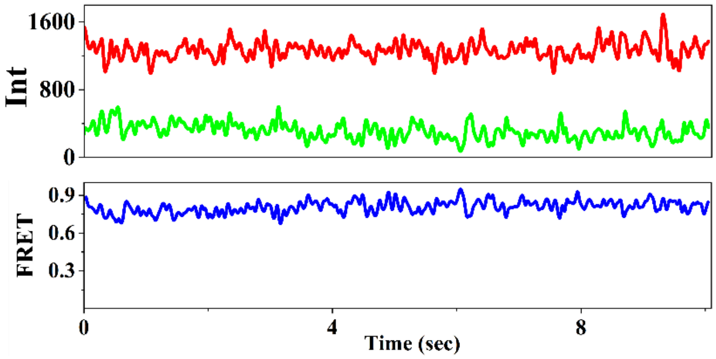


**Figure S5** Representative smFRET trace of donor and acceptor intensity versus time and FRET Efficiency versus time of a 3’ overhang fork DNA labelled at the junction

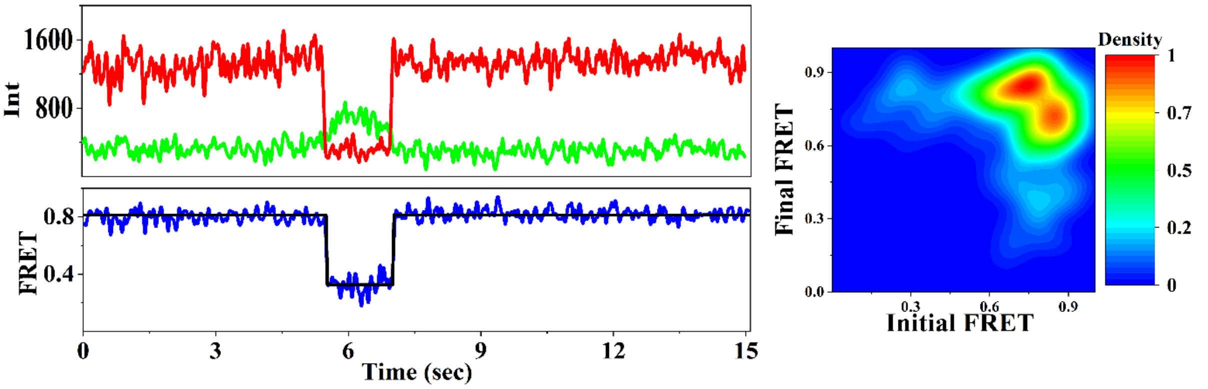


**Figure S6** Representative smFRET trace, FRET Efficiency plot versus time and TDP of a fork DNA with 5’ overhang

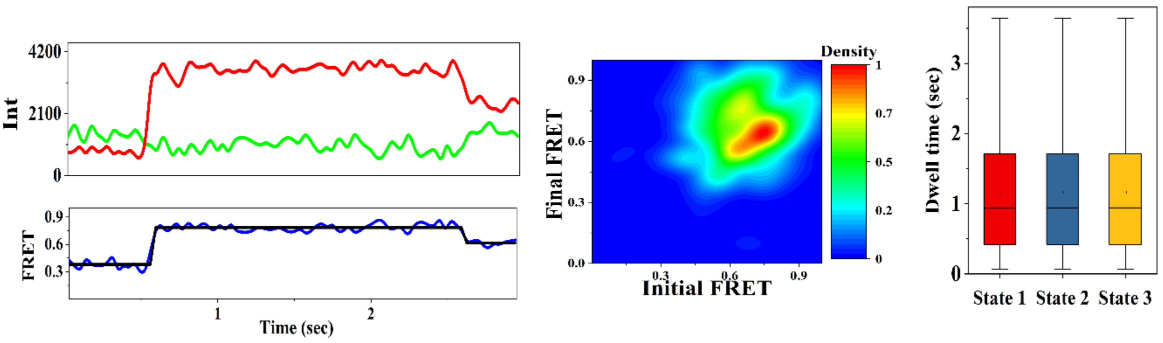


**Figure S7** Representative time trace of the donor and acceptor emission intensity and FRET Efficiency of the smFRET assay for the 3’ overhang partial fork with donor labelled at the junction and acceptor at the terminal of the nascent strand facing the junction. TDP and box plot of the dwell time distribution for the same construct. State 1 and State 2 are for the intermediate states having $E_{FRET}$ value of 0.48 and 0.61, State 3 is for the fully zipped or high FRET state.

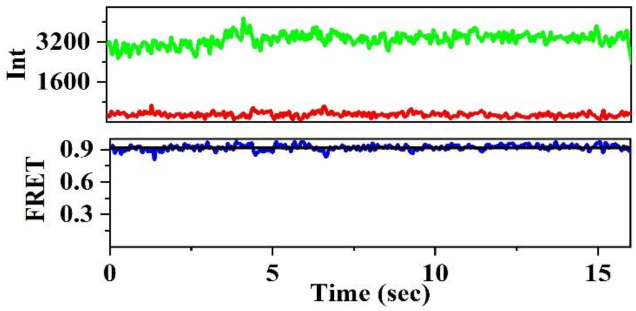


**Figure S8** Representative smFRET trace of donor and acceptor intensity versus time and FRET Efficiency versus time of a 5’ overhang fork DNA, where donor is labelled at the leading strand facing the junction and acceptor is labelled at junction of the template strand

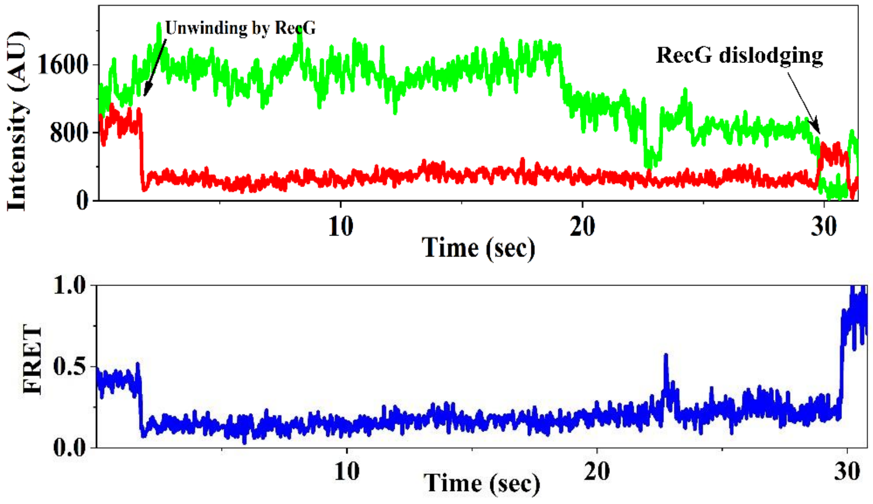


**Figure S9** Representative smFRET trace of a long movie where the blocked fork, with donor labelled at the junction and acceptor labelled at the lagging daughter strand is allowed to bind with RecG. The initiation of unwinding activity of RecG has been captured, followed by it’s dislodging after a long interval of time.
